# On-farm hatching and contact with adult hen post hatch induce sex-dependent effects on performance, health and robustness in broiler chickens

**DOI:** 10.1101/2023.05.17.541117

**Authors:** L. A. Guilloteau, A. Bertin, S. Crochet, C. Bagnard, A. Hondelatte, L. Ravon, C. Schouler, K. Germain, A. Collin

**Affiliations:** INRAE, Université de Tours, BOA, 37380 Nouzilly, France; CNRS, IFCE, INRAE, Université de Tours, PRC, 37380 Nouzilly, France; INRAE, EASM, 17700 Surgères, France; INRAE, Université de Tours, ISP, 37380 Nouzilly, France

**Author notes:** Corresponding author: Laurence Guilloteau.

## Abstract

To improve the early perinatal conditions of broiler chicks, alternative hatching systems have been developed. On-farm hatching (OFH) with an enriched microbial and stimulating environment by the presence of an adult hen is a promising solution. Day-old chicks were allotted within five hatching and rearing conditions: OFH, conventional hatchery (CH), CH and post-hatching treatment with antibiotics (CH + AB), as well as both hatching systems with an adult hen at hatching (OFH + H, CH + H). To challenge the robustness of chickens, they were exposed on D27 to suboptimal rearing conditions by combining for 4 h transport in boxes in a new room at a lower temperature and fasting. On their return to the original room, the chicken density was increased, and birds were orally vaccinated with the Gumboro vaccine. The impacts of these conditions on hatchability, chick quality score, performance, health and robustness were determined. The OFH chick body weights (BWs) were significantly greater than those of CH chicks at hatching. Whereas there was no effect of hatching conditions, the presence of hens decreased the hatchability rate, the quality score of OFH chicks and increased mortality at hatching. Treatment of CH chicks with antibiotics (CH + AB) temporarily decreased chicken BW at D19, but the feed conversion ratio (FCR) was not modified. At D19, OFH chicks had the highest BW compared to the other groups, and the presence of hens at hatching harmed chicken BW regardless of the hatching condition and FCR. An interaction between the effect of experimental rearing conditions and chicken sex was observed later for BW. In males, the OFH chickens were the heaviest compared to the other groups at D34 but not at D56. The presence of hens negatively impacted CH chicken BW at D56. In females, there was no effect of hatching condition on the BWs at D34 and D56, and the presence of hens had a positive impact on OFH chicken BW. There was no effect of hatching conditions on health parameters. In conclusion, the OFH system was a hatching system at least equivalent to the CH system. The presence of the hen at hatching and during the chick start-up phase on performance interacted with the hatching condition and the sex of the chickens.

## Introduction

The integrated management of poultry health includes maintaining health, welfare and performance throughout the life of animals. This is an even greater challenge in a global context of reducing the risk of antimicrobial resistance. One axis in the Ecoantibio2017 plan (Ecoantibio2, 2017) concerns the development of alternatives to avoid the use of antibiotics. In this context, new poultry rearing systems are being developed, particularly for the perinatal period. In poultry, the perinatal period is a stressful period for broiler chicks, which includes the hatching phase and major physiological changes to adapt to new food resources and environments. In hatcheries, chicks hatch between 19 and 21 days of incubation. They often stay more than 12 hours in the hatcher, under optimal temperature, without light and usually without access to feed and water until placement in farm buildings. The fasting period of the chicks is further increased by the time needed for hatchery processing, transportation duration and unloading at the farm, which might last up to the first 72 h after hatching. Even though chicks can use energy reserves from their yolk sac (van der Wagt et al., 2020), these conditions induce immediate and long-lasting metabolic changes (Beauclercq et al., 2019; Foury et al., 2020), behavioural impacts by increasing fear responses (Jessen et al., 2021) and consequences on chicken development, performance and welfare (de Jong et al., 2017).

To improve the early perinatal conditions of chicks, alternative hatching systems have been developed. On-farm hatching provides the chicks with immediate access to feed and water according to their needs and avoids the exposure to stressors encountered in conventional hatcheries (van de Ven et al., 2009). Eggs incubated for 18 days are transported to the farm and placed either in trays or in the litter where they hatch. The effects of these on-farm hatching systems on broiler health, welfare and performance were recently studied under commercial or more controlled conditions and had shown effects that are not always beneficial. Total mortality and footpad dermatitis in on-farm hatched (OFH) chicks were lower compared to conventionally hatched (CH) fast-growing broiler chickens (de Jong et al., 2019; 2020; Giersberg et al., 2021; Jessen et al., 2021). However, day-old chick quality was worse and breast myopathy prevalence was higher for OFH than CH chickens (de Jong et al., 2019; Souza da Silva et al., 2021).

Chicken activity and general behaviour were little affected by the hatching system, with fast-growing OFH chickens being more fearful and less active than CH chickens (Giersberg et al., 2020). Slower-growing broiler chickens hatched in organic farms tended to express less general fearfulness than CH chickens (Jessen et al., 2021a). A positive effect on growth performance was observed during the first week of life until 21 days in OFH and CH fed at the hatchery compared to CH chickens (de Jong et al., 2020), and longer when parent flocks were young (Souza da Silva et al., 2021).

Maintaining optimal health, welfare and performance of chickens is highly dependent on the gut physiology in interaction with the microbiota and mucosal immune system (Fortun-Lamothe et al., 2023). Antibiotics have been largely used in poultry production to improve performance. Growth promotion induced by antibiotics is associated with effects on the caecal microbiome at taxonomic, metagenomic, and metabolomic levels, which might be targeted via its contribution to host-microbiota crosstalk, particularly by acting on the gut barrier function (Broom, 2018; Plata et al, 2022). However, growing concerns about the increase of antimicrobial resistance in farm animals led to changes in EU and national legislation governing the use of antibiotics as growth promoters in poultry feed, which resulted in their suppression in 2006 (Council Directive 96/22/EC; Axis 2 and measure 19 of the EcoAntibio2017 plan).

Greater attention to the environment during the chick postnatal period, especially the microbial environment, is key to optimising the gut barrier function and more broadly the health and welfare of the chickens and their performance. Naturally, chicks hatch in contact with an adult hen who is a donor of microbiota and a model of learning and maternal care (Edgar et al., 2016). Early implantation of adult microbiota into the chick digestive system accelerates the maturation of the microbiota and immune system (Volf et al., 2016; Broom & Kogut, 2018; Meijerink et al., 2020). In addition, chicks reared in the presence of their mothers are less fearful than those raised without their mothers and develop more behavioural synchrony (Perré et al., 2002), even though hen genetics has a strong effect on chick behaviour, with commercial lines being less maternal (Hewlett et al., 2019). The combination of a new hatching system like OFH with an enriched microbiota and stimulating environment from the presence of an adult hen is a possible solution for chick conditions to be improved and could contribute to poultry health and welfare and product quality.

In this study, we analysed the benefits/risks of hatching systems (conventional hatcher, on-farm hatching), with the presence of an adult hen (OFH + H, CH + H) or not (OFH and CH) on hatchability and chick quality scores. We also explored the effects of these hatching conditions and the presence of an adult hen with chicks on performance, health and robustness in suboptimal rearing conditions. The combination of CH and post-hatching treatment with antibiotics (CH + AB) was added as an experimental control group of antibiotic growth promoter use.

## Animals, Materials and methods

### Experimental design

The experimentation consisted in combining different hatching conditions, chick starting with or without hens, as well as variable rearing conditions (with or without antibiotic treatment) integrating a multifactorial challenge for all conditions (Figure 1).

**Figure 1.**
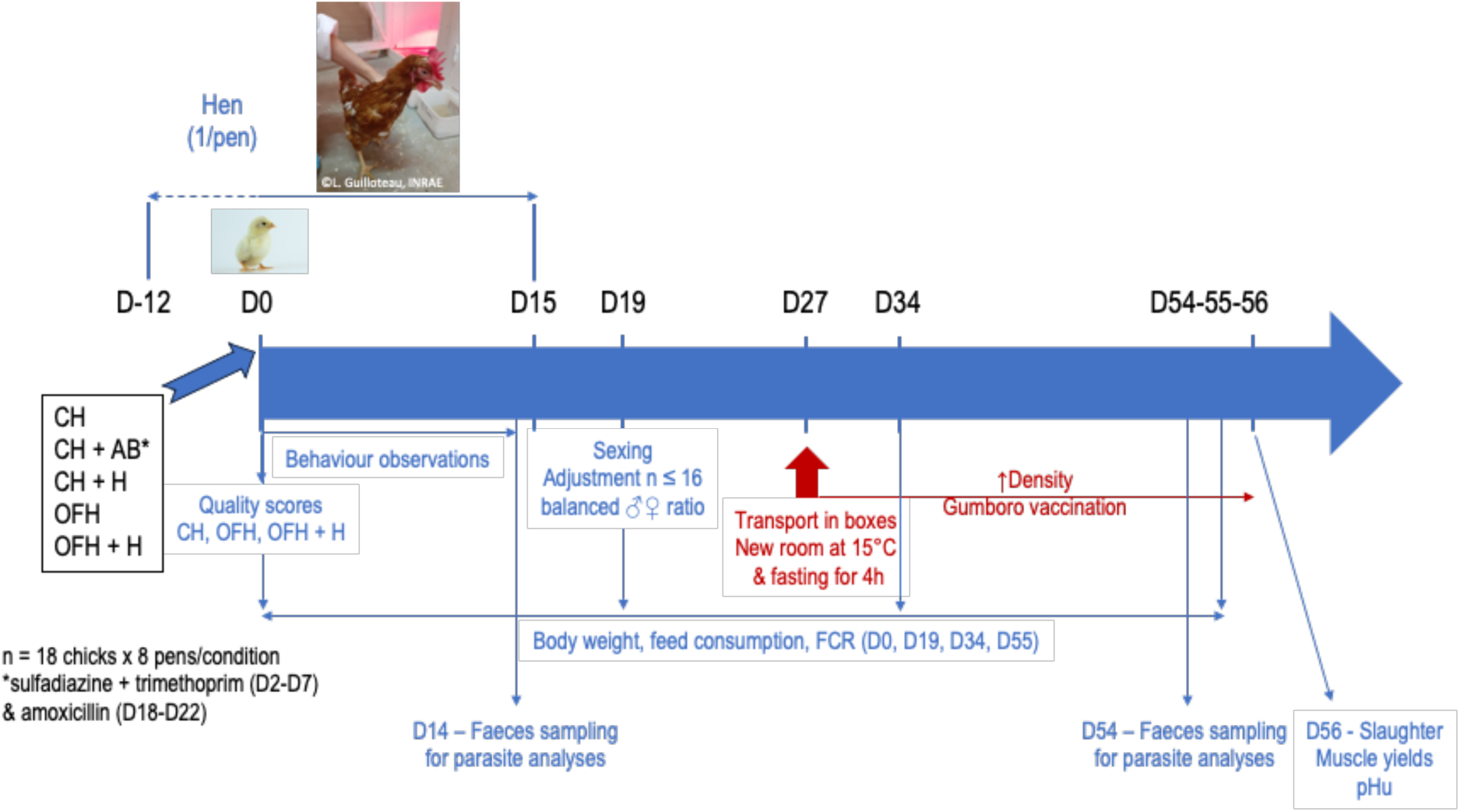
Experimental Design. Hatching conditions: conventional hatchery (CH), CH + antibiotics treatment (CH + AB), CH + hen (CH + H), on-farm hatching (OFH), OFH + hen (OFH + H).

#### Hatching conditions

Certified JA 757 18-day embryonated eggs (Galina Vendée, Essarts-en-Bocage, France) were either placed at 37.6°C with 75% relative humidity and no light in a conventional hatchery (CH) or laid directly in the litter of the pens under infrared heat lamps to allow on-farm hatching (OFH). The average temperature of the eggs in the litter was 37.9°C and under 20 h light per day until OFH chick hatching. The ambient room temperature was maintained at 25 °C with a fan heater. Day-old CH chicks were transported for one hour in a transport van before placement in pens to simulate conventional hatchery processing, which has been described to have long-term deleterious effects on fear response when combined with delayed nutrition (Hollemans et al., 2018). The time when CH chicks were placed under heat lamps in pens was considered D0 as well as for the OFH chicks already in place. Temperature under heat lamps was decreased from 35–38 °C to 31–32 °C from D0 to D3, then 29–30 °C from D4 to D6 and 26–27 °C from D7 to D13. The light cycle was 20 h light at the CH chick placement or until hatching time for OFH chick (D0), 13 h light on D1 (increased dark time to promote maternal behaviour of hens (Richard-Yris & Leboucher, 1987)), 18 h on D2 and 16 h on D3 and during the rearing period with minimum 20 lux on 80% of the lighted surface.

#### Starting period of chicks in contact with hens

Sixteen Lohmann Brown hens, acting as natural gut microbiota donors and adult presence, were obtained from a local commercial egg-laying hen farm (La cabane à Chiron, Benet, France). The hens were aged 31 weeks, vaccinated against Marek Disease Virus (MDV), Infectious Bursite Disease Virus (IBDV) and Infectious Bronchitis Virus (IBV) infections, and were sanitary controlled and declared free of *Mycoplasma gallisepticum*, *Mycoplasma synoviae, Chlamydia psittaci* and *Salmonella pullorum gallinarum.* Only *Ascaris* and *Heterakis* parasites were detected at a very low level in hen faeces.

Each hen was placed separately in a wire-latticed pen (3 m^2^) in the experimental pens described above with a nest box, perch, feed and water *ad libitum* (Figure 2A). Hens were accustomed to their new environment for 12 days, fed with a standard rearing diet for laying hens (30099G25, Arrivé Nutrition Animale, Saint-Fulgent, France) and allowed to deposit faecal and caecal materials and thus microbiota on litter. An egg was always left in the nest to encourage brooding behaviour. The room temperature was 25 °C and the artificial photoperiod was 16 h L:8 h D before egg deposition, 20 h L:4 h D during hatching and the same programme as the chicks afterwards. Two days before chick arrival or egg hatching, a wire-latticed space (101 x 50 cm) for chicks was placed in their pen (Figure 2B). Eighteen-day embryonated eggs were laid under infrared heat lamps to allow on-farm hatching (OFH) (Figure 2C). Eight hens were used for 8 groups of 18 OFH chicks, and eight hens were used for 8 groups of 18 CH chicks. On D0, day-old CH chicks were placed under the pen’s wire-latticed space, and OFH chicks were already under this space. Chicks and hens were in visual and auditory contact for a few hours. Then hens were deprived of feed and water from the morning. When lights were switched off, the hens were shut up in their nest boxes, and chicks were placed under each hen as gently as possible for 11 h without any feed and water. Chicks and hens were put physically together in the closed nest for the night to promote maternal behaviour and the acceptance of chicks (Richard-Yris & Leboucher, 1987). The nest was made of wire mesh covered with a tarpaulin and placed on shavings. The following morning, one hour before the lights were switched on, the nest-box tarpaulins were taken away to allow free access to the whole pen. The nest was present throughout the hen’s stay. Free in-access feed and water were placed under wire-latticed space for chicks (Figure 2D), not accessible for hens, and in raised troughs for hens, not accessible for chicks. Chicks could get in and out wire-latticed space as they pleased. Hens were present with chicks for two weeks, the critical period for chick start, and removed on D15. Weight and clinical examinations of the hens were recorded the day before they were installed in the pens and, on D15, when they were removed.

**Figure 2.**
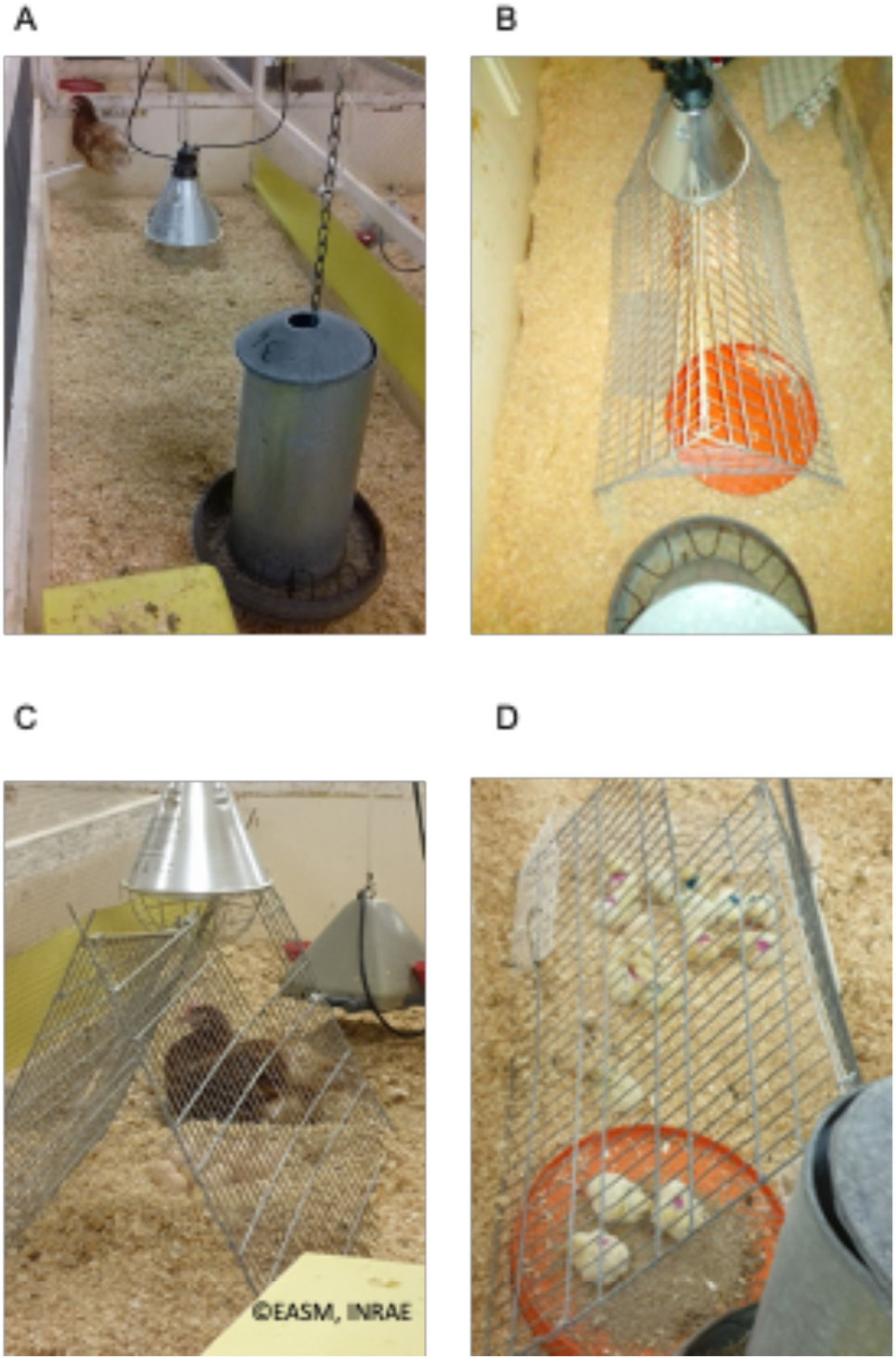
Experimental design of chick starting period in contact with hens. A. Hen wire-latticed (3 m^2^) with nest box (width 23 cm, length 35 cm, height 40 cm), perch, and free in access feed and water. B. Wire-latticed space (101 x 50 cm) for chicks within the hen pen. C. Eighteen-day embryonated eggs laid under infrared heat lamps in the chick wire-latticed space and in presence with hen. D. Chicks under the wire-latticed space with the possibility to get in and out, and to have free in access feed and water.

#### Rearing conditions

Seven hundred twenty-day-old among which 432 were from a conventional hatchery (CH) and 288 were hatched on-farm (OFH), were allocated into five groups: CH, CH + antibiotics treatment (CH + AB), CH + hen (CH + H), OFH, OFH + hen (OFH + H) (Figure 1). Each group was randomly placed in the room, repeated in eight pens (18 chicks/pen, 3 m^2^). Antibiotic treatment was only applied in chick drinking water for the CH + AB group: ADJUSOL^®^ TMP SULF Liquid (25 mg/kg sulfadiazine and 5 mg/kg trimethoprim, VIRBAC, CARROS, France) for 5 days (D2–D6) and SURAMOX 50 (400 mg/10 kg, i.e. 20 mg/kg amoxicillin, VIRBAC) for 5 days (D19–D23). Sex was determined on D19 and the number of chickens was adjusted to a maximum of 16 per pen, keeping a balanced ratio between males and females. On D27, chickens were exposed for 4h transport in boxes to a new room at a lower temperature (15 °C instead of 25 °C) and feed deprivation. On their return to the original room, the pen size was reduced from 3 m^2^ to 1.5 m^2^ to increase chicken density, and birds were orally vaccinated with the live Gumboro vaccine in drinking water (HIPRAGUMBORO^®^ -G97, HIPRA FRANCE, Saint-Herblain, France). These conditions are stress factors that chickens may encounter on farms; the objective was to expose chickens to suboptimal rearing conditions. Chickens had ad libitum access to water and to feed without any anticoccidial drugs. They were fed with a standard starter diet (raw energy = 4462 kcal/kg, crude protein = 23.91%) until D19, then a grower diet from D20 to D34 (4527 kcal/kg, crude protein = 20.51%) and a finisher diet from D35 to D56 (4600 kcal/kg, crude protein = 19.98%). A wire mesh platform and a perch were used for environmental enrichment.

### Chick quality scores

Chick quality scores were determined at placement in the pen for CH chicks (D0), corresponding to 21 days of incubation for OFH chicks, on 24 to 25 chicks from the three treatments: CH (at the entrance into the pens), OFH and OFH + H (after hatching within their pen). They were macroscopically defined according to the grid of Tona (Tona et al., 2003) and modified by adding several other parameters (Guinebretière et al., 2022). Briefly, the chicks were scored on a total score of 110, including scores of posture (on 5), down (on 5), legs (on 6), red dot on the beak (on 10), grouped into an “*appearance*” score (on 26); activity (on 6), eyes (on 16), leg joint inflammation (on 5) and leg dehydration (on 5) were grouped into a “*tiredness”* score (32), and finally, retracted yolk (on 12), navel (on 12), remaining membrane (on 12), and remaining yolk (on 16) were grouped in an “*abdomen”* score (on 52).

### Behavioural observations

The scan sampling method was used to follow the behaviour of hens and chicks on days 2, 5, 6, 7, 8, 9, 12, 13 and 14 with the following repertoire: resting (the hen is lying or standing still, eyes closed and without chicks), maintenance (preening, scratching, stretching), feeding behaviour (the hen is eating or drinking), locomotion, exploration (the hen is scratching or pecking at the ground or the environment), observation (the hen is observing the environment with neck movements), maternal behaviour (the hen is making food offering to the chicks, the hen is expressing maternal calls, the hen is brooding the chicks by lying down and spreading her wings), fear behaviour (the hen is flying or running from the experimenter, freezing, alert), agonistic behaviour (the hen is chasing the chicks, the hen is pecking the chicks, others (punctual behaviours like vocalisations). To characterise hens’ behaviour towards the chicks, each hen was categorised according to the frequencies of agonistic or maternal behaviours. We defined three categories: 1) maternal (M): the hens expressed only maternal behaviours towards the chicks; 2) tolerant (T): the hens expressed both maternal and agonistic behaviours towards the chicks or less than 5% of scans with maternal behaviour; 3) aggressive (A): the hens rejected the chicks and expressed only agonistic behaviour towards them.

To evaluate the proximity between chicks and hens, the experimenter also recorded the localisation of four chicks randomly tagged at D0 per pen and the hen within the pen. To that end, the pen was virtually divided into four zones (Figure 3). The observations were conducted between 10 AM and noon and between 3 and 5 PM by the same experimenter. The experimenter walked slowly in front of each pen and recorded the behaviour of the hen and the localisation of the four tagged chicks every eight minutes (approximately), with a total of 10 scans per hen per day and 177 scans per hen for the whole period of observation.

**Figure 3.**
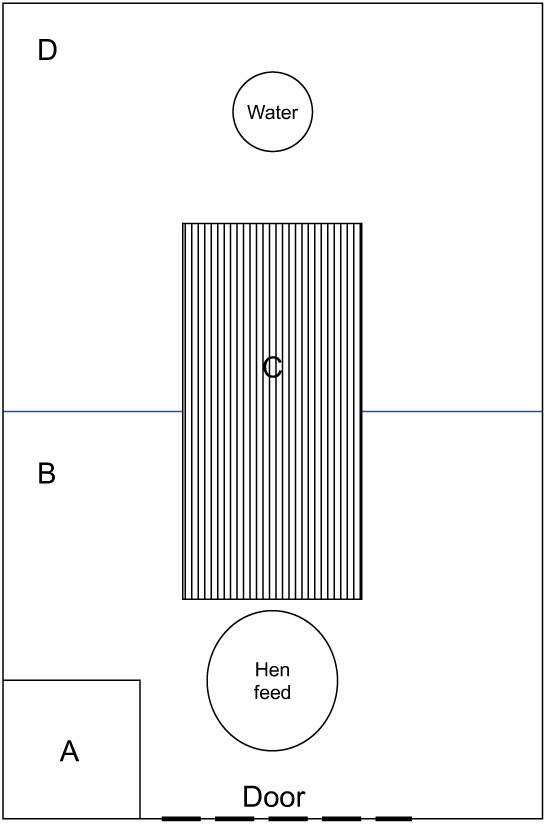
Schematic representation of the pen (3m^2^) with the zones used to locate the four tagged chicks and the hen during behavioural observations; A: the nest (23 cm wide x 35 cm long x 40 cm high), B and D: two halves of the pen and C: the wire-latticed space for the chicks (101 × 50 cm).

### Performance

Body weight (BW) was measured at D0, D19, D34 and D55. Feed consumption was measured in each pen for the periods between D0–D19, D19–D34 and D34–D55, and then used to calculate the feed conversion ratio (FCR) as the feed consumption-to-BW gain ratio per pen during both periods and the entire rearing period. At D56, 16 identified males per group were slaughtered, and *pectoralis major* and *pectoralis minor* (breast) muscles were weighed to calculate their yields relative to BW and ultimate pH. Ultimate pH was measured as the pectoralis major pH 24 hours after slaughter.

### Health parameters

Droppings deposited on pen litter were collected on D14 and D54 and analysed for parasite detection (*Coccidia*, *Ascaris* and *Heterakis*). Five grams of droppings were homogenised in 70 mL of flotation solution (0.36% of sodium chloride). The mixture was then filtered and pressed through a tea strainer (small mesh) to extract as much of the liquid part as possible. A homogeneous sample was deposited into a McMaster cell counter, and after 5 min of rest, the oocysts and nematode eggs were counted, and their number was expressed per gram of droppings (OPG). Health disorders, mortality and causes of death were registered during the experiment.

### Statistical analyses

Hatching rates between hatchery and on-farm hatchings were compared using chi-squared tests. Chick quality parameters were analysed by a non-parametric Kruskal-Wallis test, considering the treatment (CH, OFH and OFH + H), followed by Mann-Whitney post hoc tests. A 2-way ANOVA was then carried out to test the effects of the experimental group, the sex and their interaction on performance. The statistical model used was then: Yij = µ + ai + bj + abij + eij where Yij is the dependent variable, µ the overall mean, ai the Experimental group (CH, CH + AB, CH + H, OFH, OFH + H), bj the Sex effect, abij the two-by-two interaction and eij the residual error term. When there was an interaction between variables, a Fisher (LSD) test was used to determine the statistical significance of the difference. Differences were considered significant when p-values < 0.05 and a tendency for 0.05 < p < 0.1. Analyses were performed using XLSTAT software (version 2015, Addinsoft, Paris, France).

Behavioural data did not meet the assumption of normality and homogeneity of variances. Non-parametric Mann-Whitney U-tests were used on the mean percentage of scans per behavioural category to compare the behaviour of hens in contact with CH chicks to the hens in contact with OFH chicks. To compare the proximity of CH and OFH chicks towards the hen, Mann-Whitney U tests were conducted on the mean number of tagged chicks located in the same area of the pen as the hen over the 177 scans recorded per hen.

## Results

### Hatchability and chick quality

#### Hatchability

For conventional hatchers, 97.7% of CH fertile eggs hatched at E21 and 97.2% ± 4.2% of OFH fertile eggs hatched at E21 in pens. The presence of hens had a significant impact on the OFH condition (p = 0.034). In the presence of hens, 86.8% ± 11.9% of OFH + H chicks hatched at E21. Unhatched eggs were mainly found in the pens with aggressive hens (9/11) or in the OFH pens next to those with aggressive hens (4/4). No mortality of CH chicks or OFH chicks was observed at hatching, whereas 5.6% ± 5.9% (from 0 to 16.7% according to the pen) OFH + H chicks died or were removed at hatching (n = 10) due to three hens’ aggressiveness or another reason. Only 3.6% (2/56) of chicks had residual yolk sacs at the age of 20 days (one CH and one CH + AB) and no yolk residue was found at 56 days.

#### Quality scores of chicks

No difference was shown due to the hatching conditions (p > 0.05) on the total quality scores, with good scores in the three groups considered (OFH: 96.2 ± 1.5, CH: 97.3 ± 1.5; CH+H: 95.1 ± 1.7). However, the subtotal score of the appearance was impacted by treatment whereas the subtotal scores for tiredness and abdomens of the chicks were unaffected by treatment (p > 0.05, data not shown). Indeed, whereas the subtotal score for appearance was not different between CH chicks or OFH chicks, it was deteriorated by the presence of the hen within the hatching pen in OFH + H compared to OFH chicks (p = 0. 01) (Figure 4).

**Figure 4.**
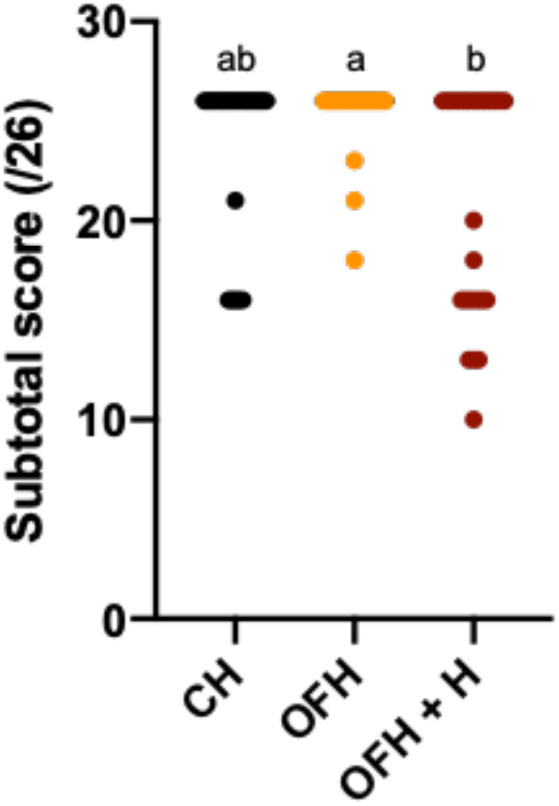
Chick appearance subtotal score at the placement in the pen according to hatching conditions; appearance scores noted on 26 included scores of posture (on 5), down (on 5), legs (on 6), and a red dot on the beak (on 10); n = 24 to 25 chicks/hatching condition; conventional hatchery (CH), on-farm hatching (OFH), OFH + hen (OFH + H)

### Behavioural observations

Because 3 hens (1 OFH + H and 2 CH + H) were very aggressive and injured their chicks, they were removed from the pens (the following day after the overnight physical contact with chicks, when they had access to the whole pen even if the chicks had access to their own space) and the later behavioural analysis. However, the chicks were kept in the analysis as they were in contact with their hen during hatching and with the microbiota the hen deposited in the pen. There was no significant difference in the behaviour of the hens, regardless of the hatching condition of chicks, except for the frequency of the behaviour “*observe*”; OFH hens tended to observe their environment less than CH hens (Additional file: Table S1).

**Table S1.** Behaviour of hens according to the chick hatching conditions.

| Hen behaviour | Hatching conditions |  | P-value |
| --- | --- | --- | --- |
|  | CH | OFH |  |
| Agonistic | 2.54 ± 3.74 | 1.37 ± 0.72 | 0.550 |
| Rest/Comfort | 17.72 ± 7.16 | 31.16 ± 22.74 | 0.181 |
| Fear | 7.07 ± 3.39 | 4.92 ± 1.95 | 0.384 |
| Feeding | 18.10 ± 4.52 | 19.45 ± 11.56 | 0.731 |
| Locomotion | 6.78 ± 4.12 | 3.39 ± 2.95 | 0.146 |
| Observation | 17.53 ± 7.45 | 9.52 ± 4.76 | **0.045** |
| Exploration | 22.62 ± 7.62 | 19.77 ± 10.78 | 0.656 |
| Maternal | 1.32 ± 1.94 | 3.39 ± 7.98 | 0.732 |
| Others | 6.32 ± 2.31 | 7.02 ± 7.02 | 0.470 |

CH = conventional hatchery (n = 6); OFH = hatching on-farm (n = 7)
Behaviour observations (mean ± SD of scan percentage over 9 days)
p-value < 0.05 = significant difference between hatching conditions (Mann-Whitney U-test)

Hens’ behaviour towards the chicks was categorised according to the frequencies of agonistic or maternal behaviours. Two hens were defined as maternal, six were tolerant, and five were aggressive among the 13 hens analysed (Table 1).

**Table 1.** Classification of hen according to the frequencies of maternal or agonistic behaviours expressed towards chicks.

| Hatching conditions | Hen behaviours |  | Category |
| --- | --- | --- | --- |
|  | Agonistic | Maternal |  |
| CH1 | 7.91 ± 0.27 | 0 | A |
| CH2 | 0 | 0.57 ± 0.07 | T |
| CH3 | 0.56 ± 0.07 | 0.56 ± 0.07 | T |
| CH4 | 0 | 5.08 ± 0.22 | M |
| CH5 | 0 | 1.69 ± 0.13 | T |
| CH6 | 6.78 ± 0.25 | 0 | A |
| OFH1 | 1.13 ± 0.11 | 0 | A |
| OFH2 | 0 | 21.47 ± 0.41 | M |
| OFH3 | 1.69 ± 0.13 | 0.56 ± 0.07 | T |
| OFH4 | 1.69 ± 0.13 | 0.56 ± 0.07 | T |
| OFH5 | 1.13 ± 0.11 | 1.13 ± 0.11 | T |
| OFH6 | 1.69 ± 0.13 | 0 | A |
| OFH7 | 2.26 ± 0.15 | 0 | A |

CH = conventional hatchery; OFH = on-farm hatching
Behaviour observations (mean ± SD of scan percentages over 9 days)
A = Aggressive
T = Tolerant
M = Maternal

The mean number of chicks observed in the same area as the hen did not differ significantly between CH (0.42 ± 0.14, n = 6) and OFH (0.39 ± 0.21; n = 7) chicks (p > 0.05).

### Performance

Hatching conditions significantly influenced chick BW from hatching to slaughter age. The OFH chick BW was significantly greater than that of all CH chicks at hatching, whether hens were present or not (p ≤ 0.002, Figure 5). A sex effect was observed from D19 onwards; male chicken BWs were greater than those of females (males: 503 ± 46g, females: 469 ± 37g, p = 0.0001). Treatment of CH chicks with antibiotics temporarily decreased chicken BW at D19 (p = 0.035) (Figure 5) due to a decrease in weight gain in females (Table 2) compared to CH chickens, while feed intake (data not shown) and FCR were not different (Table 2). At D19, OFH chickens had the best BW compared to all other groups of chicks (p ≤ 0.0003) (Figure 5) and the best weight gained per chicken (Table 2). At this time, the presence of hens at hatching with CH and OFH chicks had a remnant negative impact on chicken BW regardless of the hatching condition (p < 0.0001), as well as on weight gain and FCR for the period D1-D19 (Table 2). Both the feed intake per chicken (CH: 624 ± 12g^a^, CH + AB: 600 ± 27g^ab^, CH + H: 603 ± 25g^bc^, OFH: 652 ± 33^a^, OFH + H: 615 ± 34^c^, p = 0.001) and the weight gained per chicken (Table 2) decreased compared to the other groups, and the FCR increased (Table 2). An interaction between the effect of the experimental group and chicken sex on BW was observed later at D34 (p = 0.012) and D56 (p = 0.022) on BW, even though the FCR was not affected (Table 2). At D34, a week after the challenge, the OFH male chickens were the heaviest compared to the other groups (p ≤ 0.033) and the best weight gain (Table 2). The presence of hens at hatching harmed chicken BW (p ≤ 0.0004), regardless of the hatching condition (Figure 6A) and the FCR was not affected (Table 2). In females, there was no effect of hatching condition or presence of hens on the BW at D34 (Figure 6A). At slaughter age (D56), there was no effect of hatching condition on the male chicken BW, but the presence of hens at hatching harmed CH chicken BW (p = 0.0008) (Figure 6B) and weight gain for the period D34 – D56 (Table 2). There was a pen effect in CH + H (p = 0.016) and OFH + H chickens (p = 0.001), the pen with the lightest CH + H males was in the presence of an aggressive hen, and the heaviest OFH + H males were in a pen in the presence of a tolerant hen, but all combinations were observed (Additional file: Figure S1). In females, there was no effect of the hatching condition on the BW. The presence of hens at hatching had a positive impact on OFH female chickens compared to CH female chicken BW (p = 0. 0096), with the OFH + H chickens being the heaviest compared to the other CH female conditions (Figure 6B), and having the best weight gain for the period D34 – D56 (Table 2). There was no significant pen effect between CH + H and OFH + H female chickens (p = 0.447).

**Table 2.**
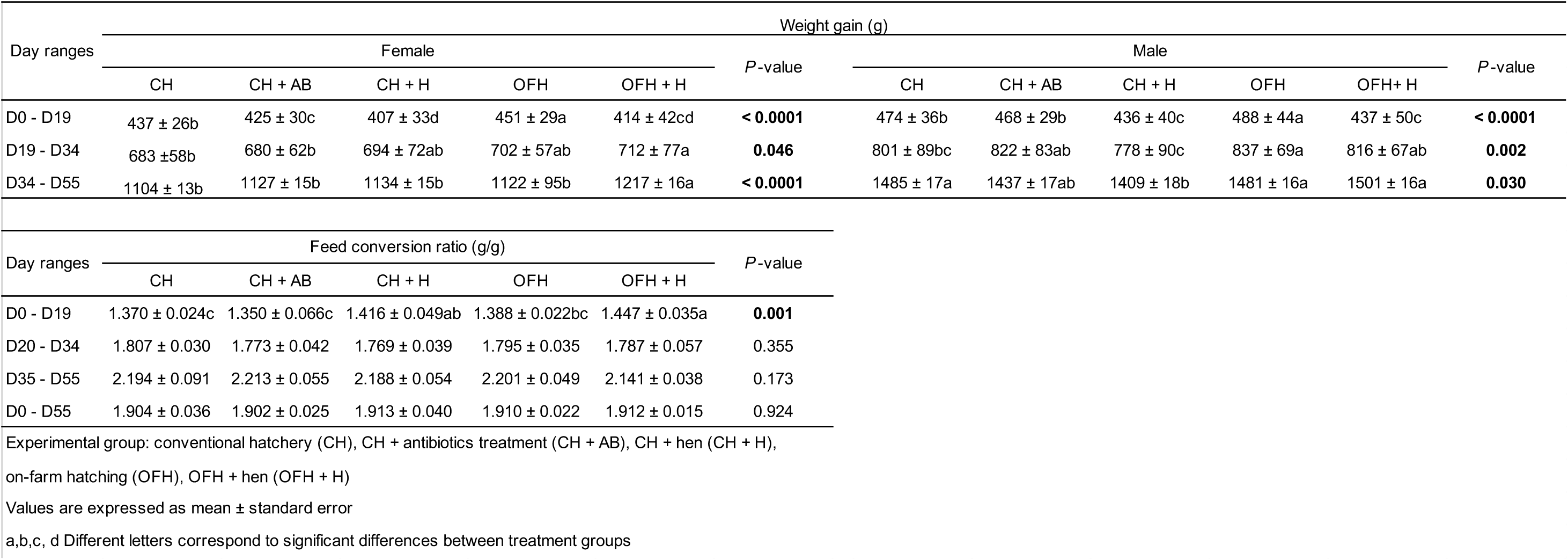
Performance according to the experimental group of chicks.

**Figure 5.**
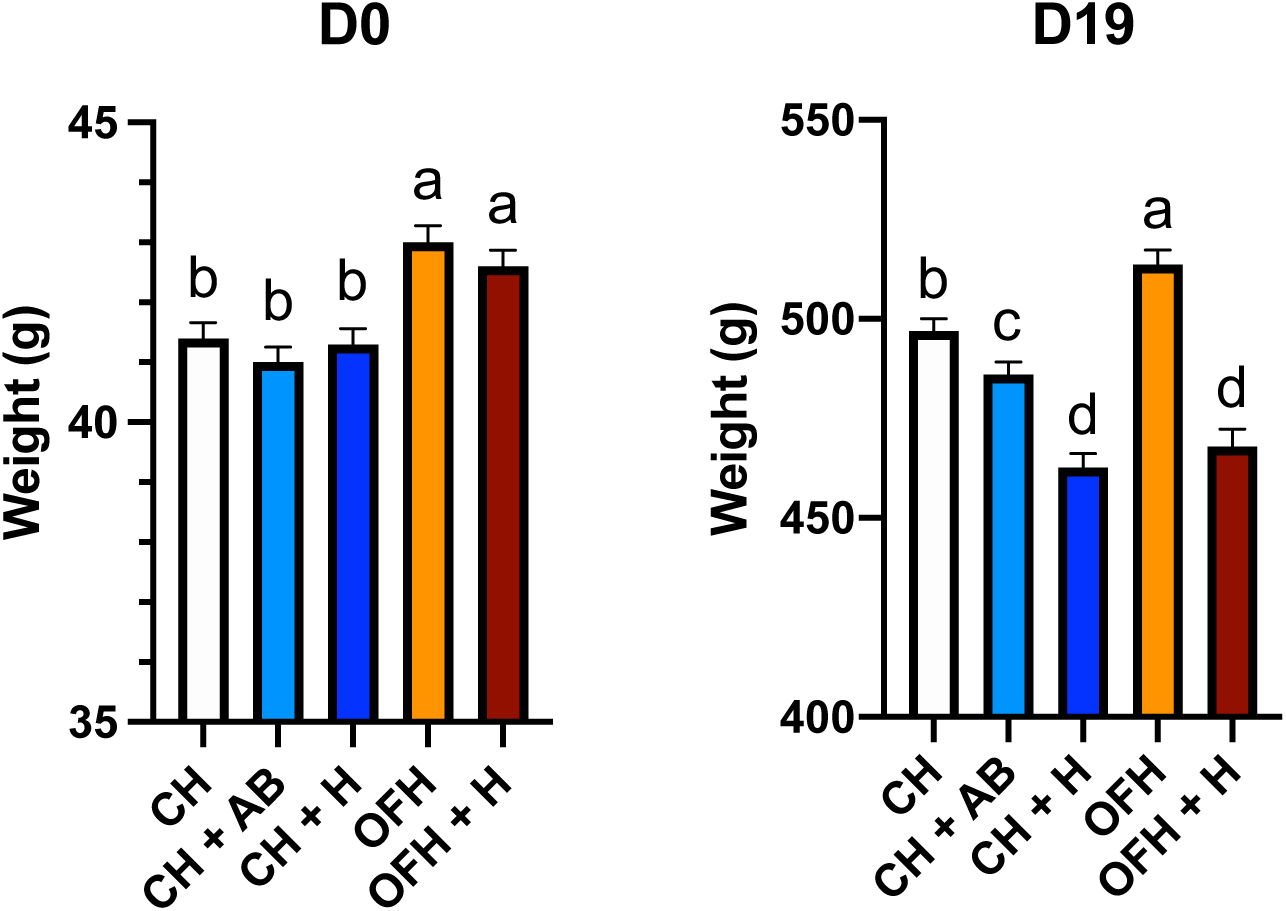
Body weight at D0 and D19 and according to the hatching conditions: conventional hatchery (CH), CH + antibiotics treatment (CH + AB), CH + hen (CH + H), on-farm hatching (OFH), OFH + hen (OFH + H); values are expressed as means ± standard error; different letters correspond to significant differences between treatment groups

**Figure 6.**
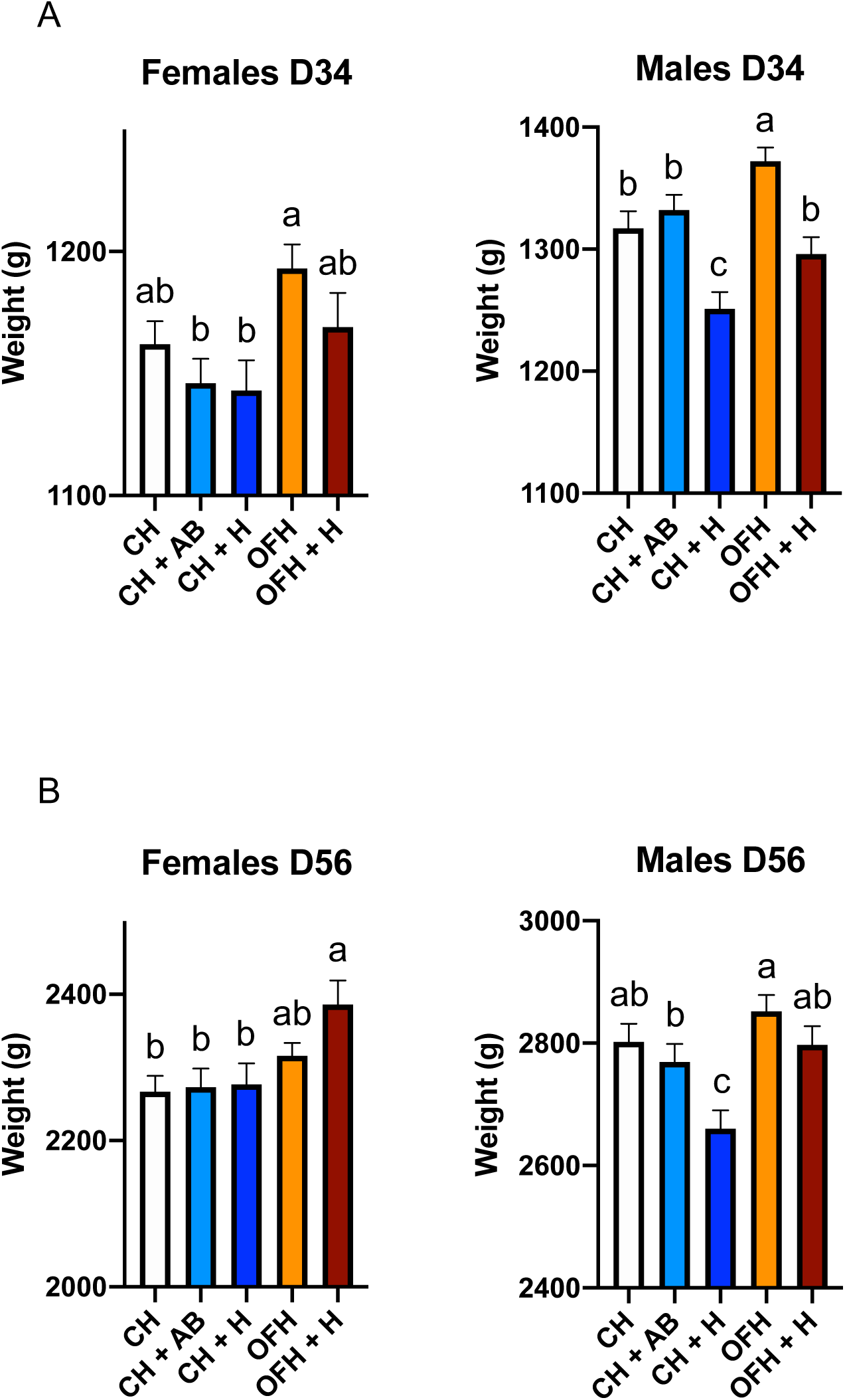
Weight at D34 (A) and D56 (B) of male and female chickens according to the hatching conditions: conventional hatchery (CH), CH + antibiotics treatment (CH + AB), CH + hen (CH + H), on-farm hatching (OFH), OFH + hen (OFH + H); values are expressed as mean ± standard error: different letters correspond to significant differences between treatment groups

**Figure S1.**
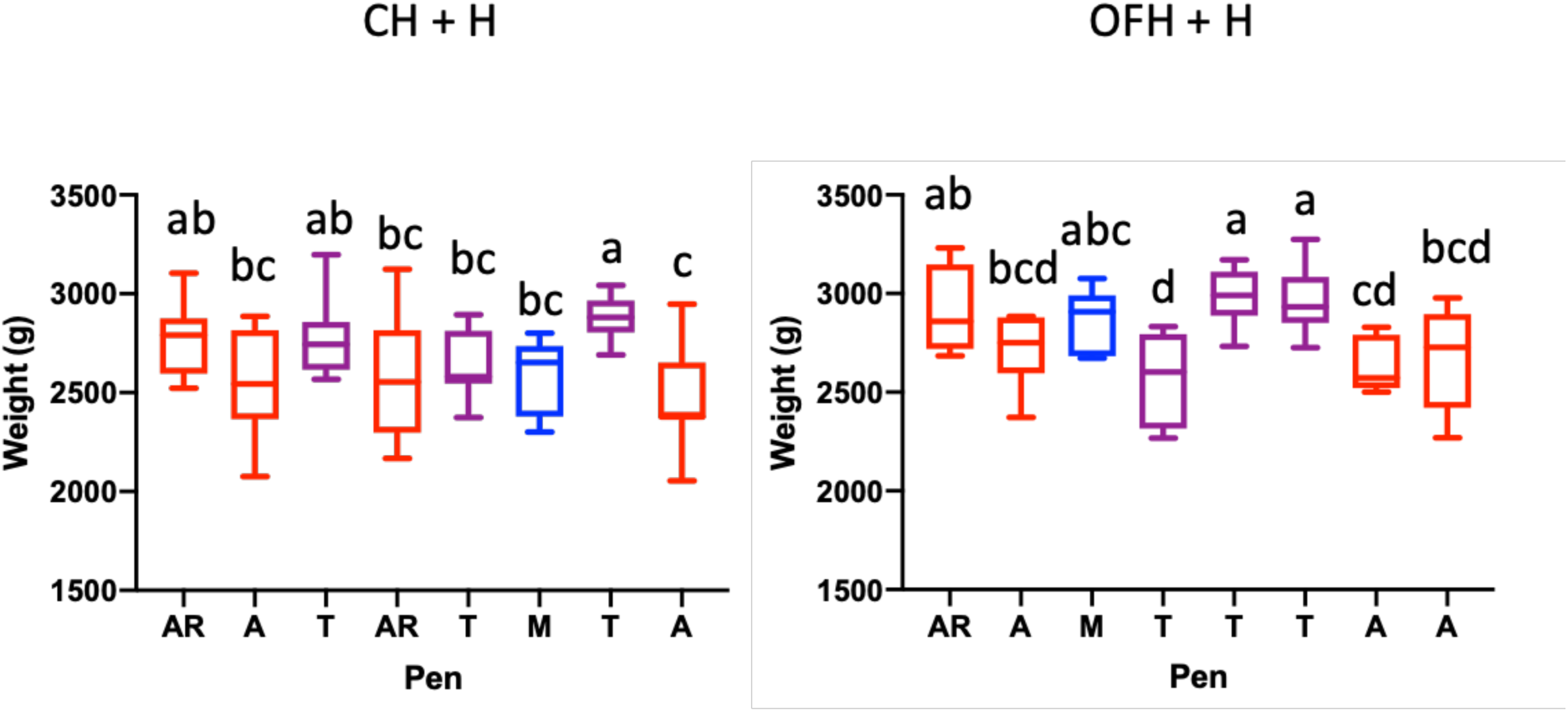
Body weight at D56 of male chickens according to the behaviour of the hen present at the starting period, M: maternal, T: tolerant, A: aggressive, AR: aggressive and removed from the pen; CH + H: chicks hatched in the hatchery and in the presence of hens; OFH + H: chicks hatched on-farm in the presence of hens; median ± SD (n ≤ 9).

Breast weight was not affected by the hatching conditions (6.99 ± 0.06, p = 0.357) and ultimate pH was not modified either (5.7 ± 0.1, p = 0.951).

### Health and robustness

*Coccidia* was detected in variable amounts in the droppings of all the pens at D54 (200–85500 OPG) without any significant effect of the hatching conditions in the presence of hen or not (p = 0.606). No clinical signs were observed during the experiment. In all hatching conditions combined, the viability rate of the chickens was 95.3%. The mortality rate during the whole experiment was 3.19% (23/720). Seventeen chicks died during the first week of life, 11 OFH + H and 5 CH + H in the presence of hens and one OFH chick for an unknown reason. Six CH chickens died during the rest of the experiment, five of which were due to heart problems (2 CH, 1 CH + AB, 2 CH + H) and one to unknown causes (CH + H). Eleven chicks were additionally eliminated after hatching in pens in the presence of hens (4 at D1, 4 at D2, and 1 at D4) and two later (D33 and D55) for morphological reasons.

## Discussion

New hatching systems are being developed in Europe, and the enrichment of the rearing environment is also in full development, notably by optimising the microbial environment of the chicks to limit the use of antibiotics. In this study, we analysed the benefits/risks of hatching systems (OFH and CH, treated with antibiotics or not) and of the presence of an adult hen or not on hatchability, chick quality score, performance, health and robustness.

### Hatching conditions

The hatching conditions compared within the present study concerned a combination of environmental parameters diverging for both hatching conditions (hatcher or on-farm), from the light regimen to the hatching temperature and the relative humidity, and the egg position. Additionally, there was a partial contact with the litter through the floor-hatching device compared to the hatcher crate. The BW of OFH-certified JA757 chicks was significantly greater than that of CH chicks at hatching, even though the hatchability rate and the quality score of chicks were comparable between the two conditions, and no mortality was reported. These results agree with other studies performed on larger number of fast-growing broilers in terms of BW, but not in terms of chick quality, which was lower in OFH chicks than in CH chicks (de Jong et al., 2020; Souza da Silva et al., 2021). In OFH-slow-growing organic broilers, BW was also reported greater, as well as the hatchability, and of lesser chick quality than that of CH chicks at hatching (Jessen et al., 2021a; Jessen et., 2021b). However, in our study, there was no effect of hatching conditions, but the presence of hens decreased the hatchability rate, the appearance quality score of OFH chicks and increased mortality at hatching. The negative effect on these indicators could be linked to the very few hens expressing a clear maternal behaviour towards the chicks (n = 2/16); some of them even showed agonistic behaviour. However, this genetic line was chosen because the studied practice could favour the possibility to use culled hens in breeding, and because of their rather tolerant behaviour, it may be possible to optimize their brooding behaviour. Improvements could be obtained by carrying it out in a season with days with greater light amplitudes (spring) to facilitate brooding behaviour, which was not the case in this study (winter), and by selecting hens with brooding behaviour to facilitate maternal behaviour (Shimmura et al., 2010). Light color and intensity are also known to influence social interaction between hens, and tuning both the color and the intensity could be a management strategy to decrease aggressive behaviour such as pecking but whose effects vary according to age, genetics and activities (Du et al, 2022). In addition, in our experimental design, the chicks had to feed under the wire-lattice space, which was not accessible to the hen. As they obtained both food and warmth under this space, the hens probably did not have enough tactile stimulation from the chicks to fully express their maternal behaviour with no agonistic behaviour. Indeed, in addition to the physiological state, tactile stimulations from chicks play an important role in the expression and maintenance of maternal behaviour in hens (Richard-Yris & Leboucher, 1987).

### Starting period

Hatching conditions and the presence of hens for 15 days after placement significantly influenced chick performance during the starting period. At D19, OFH chicks had the highest BW compared to the other groups. No significant differences were observed in the behaviour of hens present with OFH and CH chicks, except for OFH hens, which were found to observe their environment less than CH hens. With our small sample size, this result could be explained by the behaviour of one OFH hen, which spent much of the time resting. The CH and OFH chicks did not differ in their proximity towards the hen. The mean number of chicks observed in the same area as the hen was very low (less than 1 chick), indicating that they were rarely in contact with the hen. However, chick performance was affected by the presence of the hens, including lower feed intake and consequently lower weight gain and higher FCR. This could be explained by the agonistic behaviour of some hens towards chicks, the attempt of the hens to eat the chick feed and the stress that this may have caused the chicks.

Treatment of CH chicks with antibiotics, assessed as growth promoters, temporarily decreased chicken BW at D19, but FCR was not modified. This effect was not observed later, but growth promotion was not observed in CH chicks treated with antibiotics. This result is not in agreement with the use of antibiotics as growth promoters in farm animals, but the relative lack of published data on chicken performance limits knowledge of the actual effects of antibiotics on animal performance (Kumar et al., 2018; Broom, 2018; Plata et al, 2022). Their effects also result from their interaction with the microbiota and the variables chosen in the experimental studies.

The effects observed in farms are dependent on the sanitary conditions present, which are different from the much more controlled sanitary conditions in the experimental studies and may contribute to different effects of treatment with antibiotics.

### Growth period

An interaction between the effect of hatching conditions and chicken sex was observed on BW after the challenge on D27. In males, the OFH chicken group was the heaviest compared to the other groups at D34 but not at D56. These results are consistent with a previous study that observed the beneficial effects of OFH on BW only until D21 (de Jong et al., 2020), and not until slaughter time, as reported in various studies when post-hatching feed deprivation time was at least 36 h (de Jong et al., 2017). This may reflect late compensatory growth in CH chickens that have feed deprivation after hatching. Indeed, weight gain between CH and OFH chickens was no longer different from D19 for females, and from D34 for males. Alternatively, this may also be a result of the response to the challenge experienced by the chickens at D27, including transport, exposure to low temperature, transient feed deprivation, vaccination and a change to a higher rearing density, but in fact there is no ultimate positive impact of OFH on BW at slaughter time. Moreover, in our conditions, the presence of hens eventually negatively impacted male chicken BW, but only for CH chickens at D56. In females, there was no effect of hatching conditions on the BW at D34 and D56, and the presence of hens eventually had a positive impact on OFH female chicken BW. These results were unexpected, but it is known that early stress induces sex-specific, immediate and life-long effects on the stress response, behaviour, sex hormones, and hypothalamic and blood gene expression in chickens (Madison et al., 2008; Elfwing et al., 2015; Foury et al., 2020), with the males being more reactive than the females. The results observed in this study raise questions about the consequences of hatching conditions in the presence of a hen according to the sex of the chicks. It can be assumed that male chicks developed more fear and stress responses than females when placed in the presence of a hen, and this had negative effects on their growth until slaughter age for CH chicks. For male OFH chicks, in which the effect of hen presence on their growth was only observed during the growth phase, the communication between hens and embryonated eggs before hatching (Edgard et al, 2016) and with chicks at hatching that may have a more limited effect on their growth. This could even have had negative consequences on hatchability and mortality rates, but the sex of the chicks was not recorded at that time. The presence of hens with the female OFH chicks did not affect their performance and even had a beneficial effect on their growth at slaughter age. These differences observed between treatments and chick sexes for performance are not likely explained by a difference in proximity between hens and chicks, which was low in this experiment.

### Health and Robustness

There were no effects of hatching conditions on health parameters (parasitic load, clinical signs, rate of mortality), even after exposure of chickens during their growth phase to an environmental and vaccine challenge. One limitation of the experiment is that it does not reflect the farm environment which may include an accumulation of stressors in a more complex health environment. An infectious challenge could test the potential benefits of these rearing conditions. However, the challenge used in this study could have accentuated the differences in the effects of hatching conditions on performance parameters between males and females, but we did not perform the unchallenged rearing conditions to assert this. The implantation of adult microbiota into the chick digestive system by the presence of hens should be nevertheless beneficial for the maturation of the chick microbiota and gut immune system and still needs to be assessed.

Altogether, on-farm hatching of certified broilers was a hatching system at least equivalent to the hatchery hatching system in this study. The possibility of adding the presence of a hen at chick start-up remains tricky. The health status of the hens was controlled to ensure that no pathogens were transmitted to the chicks. However, the presence of hens, categorised according to their behaviour, revealed deleterious effects on hatching rate, the appearance quality score and hatching mortality. So, the health status and behaviour of the hens are essential to ensure the health status and welfare of the chicks. Moreover, the effects of the hens’ presence at hatching and during the chick start-up phase on performance interacted with the hatching condition and the sex of the chickens. To better study hen-egg/chick interaction, the sex effect could be better characterized by in ovo sexing. Further studies should be done to assess the effects of these hatching and chick-starting conditions, in the presence or absence of hens, on the implantation and maturation of the chicks’ gut microbiota and mucosal immunity. New devices enabling interactions between hens and chicks should also be tested.

## Ethics approval

All experimental procedures were approved by the Ethics Committee COMETHEA POITOU-CHARENTES n°84 (APAFIS#24474-2020021816237418 v3) and carried out following current European legislation (EU Directive 2010/63/EU).

## Author contributions

LAG, AB, CS, KG and AC designed the study with the help of CB. LAG, CB, AC and CS performed the experiment with the technical help of SC for the organisation of the experiment and AH for parasitic analyses. CB and LR collected the performance and health parameters. LAG analysed data with the help of AB and CB for the behaviour data. LAG, AB and CB wrote the paper with the help of KG and AC. All the authors reviewed and approved the manuscript.

## Acknowledgements

We are grateful to all the members of the RIMEL network whose shared thinking made the design of this study possible. We thank the staff of the poultry alternative breeding experimental unit (EASM, INRAE, 17700 Surgères, France, DOI: 10.15454/1.5572418326133655E12) for the development of the experimental set-up and the conduct of the experimentation. We are very grateful to the staff of the MOQA team (INRAE, 37380 Nouzilly, France) for their help during the experimentation and Sandrine Grasteau for her advice on statistical analyses. The manuscript has been professionally proofread.

## Data and model availability statement

The datasets used during the current study are available on line: https://doi.org/10.57745/6INVYL

